# Oxidative inhibition of PTEN links ROS signaling to regeneration and homeostasis

**DOI:** 10.64898/2026.07.30.741702

**Authors:** Iván Sopena-Majós, José Esteban-Collado, Joan Vallhonrat-Pinell, Lidia Pérez, Hugo Stocker, Montserrat Corominas, Florenci Serras

## Abstract

Reactive oxygen species (ROS) are early signals of tissue damage, but how they engage growth-promoting pathways in vivo is unclear. Here we show that ROS promote regeneration by inhibiting PTEN, thereby activating PI3K–Akt signaling in *Drosophila melanogaster*. We identify cysteine 79 as an essential residue for redox regulation of PTEN, demonstrating that a ROS-insensitive PTEN mutant uncouples Akt activation from oxidative stress. This disruption impairs regenerative growth in wing imaginal epithelia and stem cell proliferation in the adult gut. Mechanistically, Akt activation links ROS signaling to p38 MAPK-dependent regeneration responses. Together, our findings define a conserved ROS–PTEN–Akt signaling axis that integrates oxidative damage with regenerative growth, establishing PTEN as a key redox-sensitive regulator of tissue repair.

## Introduction

Tissue regeneration is a finely orchestrated biological process that restores structure and function following injury. Among the many signals that initiate and control this complex response, reactive oxygen species (ROS) have emerged as crucial early mediators (Diwanji & Bergmann, 2017, 2018; Fox et al., 2020; Niethammer, 2018; Serras, 2016). Once considered merely cytotoxic by-products of cellular metabolism, ROS are now recognized as dynamic signaling molecules that initiate and modulate regenerative responses across multiple organisms (Ferreira et al., 2016; Fogarty et al., 2016; Gauron et al., 2013; Jaenen et al., 2021; Khan et al., 2017; LeBert et al., 2018; Love et al., 2013; Rampon et al., 2018; Sena & Chandel, 2012; Wenger et al., 2014).

One such pathway is the phosphoinositide 3-kinase (PI3K)-Akt pathway, a central regulator of cell survival and proliferation (Manning & Toker, 2017), which is activated during regeneration (Esteban-Collado et al., 2021; Santabárbara-Ruiz et al., 2019). Biochemical and cell culture studies have shown that ROS can oxidatively inhibit the phosphatase and tensin homolog (PTEN), an antagonist of the pathway, thereby enhancing PI3K-Akt signaling (Burge et al., 2025; Kwon et al., 2004; Leslie et al., 2003; Trinh et al., 2024). Oxidation-dependent PTEN inactivation promotes phosphatidylinositol (3,4,5)-trisphosphate (PIP₃) accumulation and Akt phosphorylation, suggesting that ROS could activate PI3K-Akt signaling during tissue damage responses. However, whether PTEN oxidation mediates ROS-dependent Akt activation during tissue regeneration in vivo remains unknown.

One downstream consequence of PI3K-Akt activation during regeneration may be the modulation of stress-activated MAPK signaling. Previous studies have shown that regeneration requires both p38 and JNK signaling (Bergantiños et al., 2010; Brock et al., 2017; Diwanji & Bergmann, 2020; Khan et al., 2017; Patel et al., 2019; Santabárbara-Ruiz et al., 2015, 2019; Smith-Bolton et al., 2009a; Sun & Irvine, 2011). The upstream MAP3K ASK1 activates these MAPKs but also promotes apoptosis (Ichijo et al., 1997; Nishida et al., 2017; Obsil & Obsilova, 2017; Tobiume et al., 2001). We previously demonstrated that ROS-induced Akt activation attenuates ASK1 activity, thereby limiting apoptosis while preserving regenerative p38 signaling (Esteban-Collado et al., 2021; Santabárbara-Ruiz et al., 2019). However, the upstream mechanism linking tissue damage-induced ROS production to activation of the Akt-ASK1-p38 regenerative signaling axis has not been established.

We hypothesized that damage-induced ROS promote regenerative signaling by transiently oxidizing and inhibiting PTEN, thereby activating the PI3K-Akt–ASK1–p38 signaling axis. To test this hypothesis, we analyzed ROS-dependent PI3K-Akt signaling in two complementary *Drosophila* epithelial regeneration models: larval wing imaginal discs, which regenerate through compensatory proliferation (Fan & Bergmann, 2008; Herrera et al., 2013; Hsu & Smith-Bolton, 2025; Repiso et al., 2013; Smith-Bolton et al., 2009b; Verghese & Su, 2016; Worley et al., 2022), and the adult midgut, where tissue repair depends on intestinal stem cells (Micchelli & Perrimon, 2006; Ohlstein & Spradling, 2006; Zeng & Hou, 2015).

Here, we show that tissue damage induces PTEN oxidization, resulting in PI3K-Akt activation in regenerating wing imaginal discs and the adult midgut. We further demonstrate that this redox-sensitive PTEN-Akt module promotes p38-dependent regenerative signaling, identifying PTEN as a redox-sensitive regulator that links oxidative stress to tissue repair.

## RESULTS

### Activated Akt rescues regeneration under nutrient-restricted conditions

We have previously shown that regenerative Akt activation depends on both ROS and nutrient availability: genetic ablation increases Akt phosphorylation, whereas nutrient deprivation or inhibition of ROS production suppresses Akt activation and impairs regeneration (Esteban-Collado et al., 2021; Santabárbara-Ruiz et al., 2019). However, whether Akt activation is sufficient to rescue regeneration under nutrient-restricted conditions has not been directly tested.

To study regeneration in the *Drosophila* wing imaginal disc, we used a genetic system that permits tissue ablation with precise spatial and temporal control (Fig. 1A). It consists of a double transactivator engineered to have two functions (Santabárbara-Ruiz et al., 2015). One component is a modified version of the LexA (hereafter *LHG*) transactivator system that can be conditionally controlled by the temperature-sensitive *Gal80^TS^* (Yagi et al., 2010). We used the wing-specific *sal^E/Pv^* enhancer to drive the expression of LHG in the cells of the central part of the wing pouch where the pro-apoptotic construct *lexO-rpr* was activated (*sal^E/Pv^-LHG lexO-rpr;* hereafter *sal^E/Pv^>rpr)*. The second component is the *Gal4/UAS* transactivator, which drives expression of the *UAS-Pdk1:UAS-Akt1* recombinant construct (hereafter *Pdk1:Akt*), in a broader wing pouch domain under the control of the *nubbin-Gal4* (hereafter *nub>*) (Fig. 1A). This transgene allows co-expression of *Akt* and *phosphoinositide-dependent kinase-1* (*Pdk1*), the gene encoding the kinase that phosphorylates and activates Akt. This construct ensures the overexpression of both genes and the activation of Akt.

**Figure 1.**
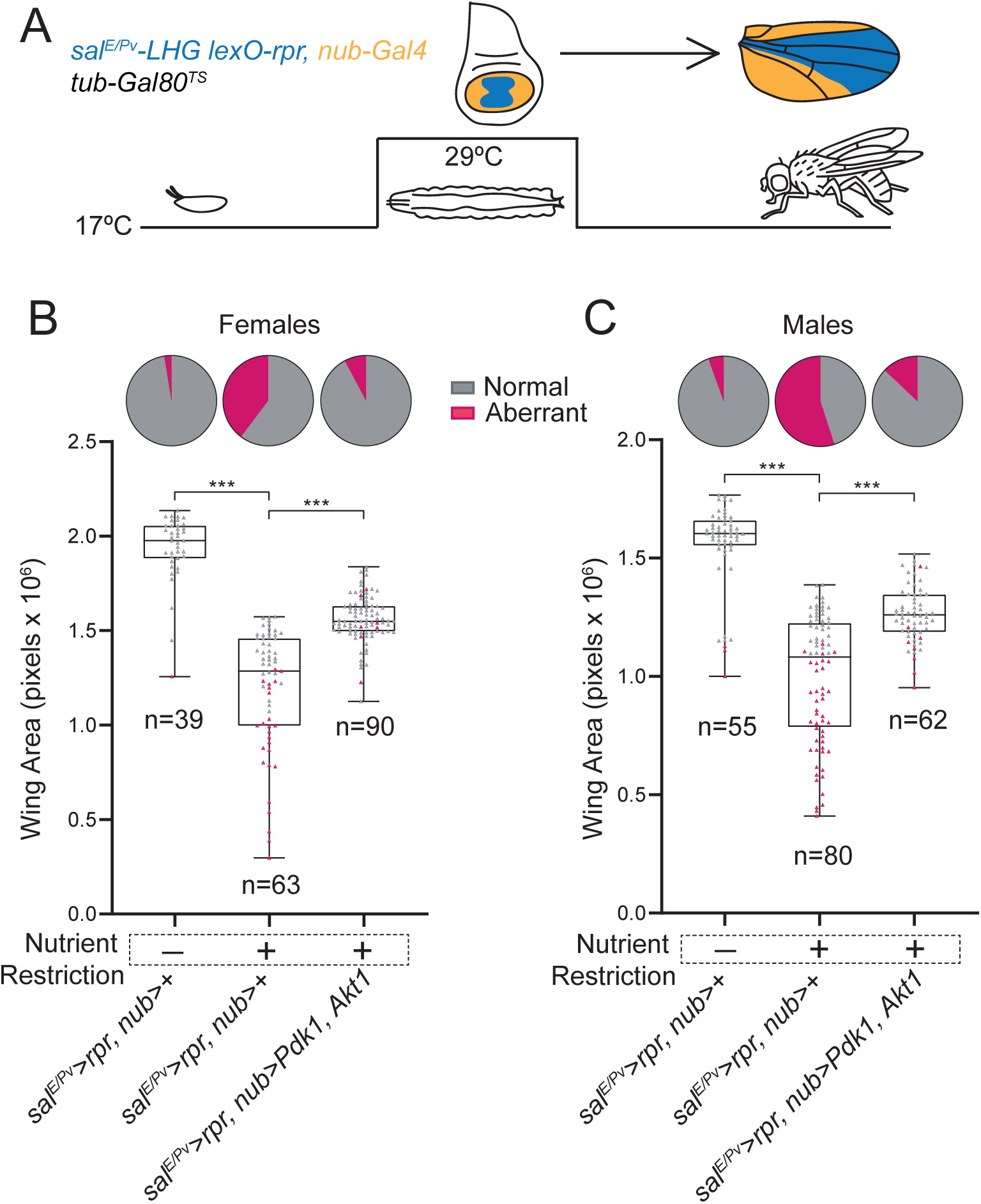
Akt activation rescues regeneration after nutrient restriction. (A) Schematic of the dual transactivation system used in the wing imaginal disc (top). Blue, apoptotic region generated by *sal^E/Pv^-LHG LexO-rpr* (*sal^E/Pv^>rpr*); orange, transgene activation driven by *nub- Gal4* (*nub>*) to express the indicated UAS-transgene. The corresponding regions in the adult wing are indicated. Experimental protocol: embryos were maintained at 17°C and 8 days after egg laying the temperature was shifted to 29°C to inactivate the Gal4 repressor Gal80, thereby inducing both *rpr*-mediated apoptosis and transgene expression. (B, C) Effects of nutrient restriction on wing regeneration. Top: Pie charts showing the frequency of regenerated wings (gray) versus non- regenerated, aberrant, wings (purple). Bottom: Box plots showing adult wing areas under standard food conditions, nutrient restriction, and nutrient restriction with Akt overexpression. Adult female and male wings were analyzed separately. Box plots show the maximum–minimum values (whiskers), the upper and lower quartiles (boxes), and the median value (horizontal line). Genotypes and sample sizes are indicated at the bottom. Statistical significance was determined by one-way ANOVA followed by Tukey’s multiple-comparisons test: ***p < 0.001.

First, we induced apoptosis (*sal^E/Pv^>rpr)* in the wing imaginal disc to stimulate regeneration in standard versus nutrient restriction conditions. To assess the regeneration capacity, we analyzed two parameters, the wing area and the vein-intervein patterning. As expected, nutrient restriction significantly reduced the size of regenerated wings and increased the frequency of aberrantly regenerated wings (Fig. 1B, C; Supplementary Fig. 1). Remarkably, while the reduction of wing size after nutrient restriction was comparable between males and females, patterning defects were more frequent in males. Under standard food, 97.4% females versus 94.6% males regenerated normal shaped wings. In contrast, nutrient-restricted conditions reduced the proportion of normally regenerated wings to 60% in females and 45% in males. This suggests that females regenerate slightly better than males, and that nutrient restriction strongly impacts the regenerative capacity, especially in males.

Next, we overexpressed *Pdk1:Akt* in a broader surrounding domain (*nub-Gal4*) to promote phospho-Akt production under nutrient-restricted conditions (Fig. 1A-C). Importantly, Akt activation significantly rescued the anomalous regeneration induced by nutrient restriction in both female and male flies (Fig. 1B, C; Supplementary Figure 1). These results demonstrate that reduced phospho-Akt availability under nutrient restriction limits regenerative capacity and establish Akt phosphorylation as a pivotal event for regeneration. The recovery after *Pdk1:Akt* activation was proportionally similar in females and males, reaching 92.2% and 87.1% of regenerated wings, respectively.

To test whether Akt phosphorylation responds directly to oxidative stress, we cultured wing imaginal discs in the presence of increasing concentrations of H₂O₂ resulting in a dose-dependent increase in Akt phosphorylation (Fig. 2A). A similar effect was observed in discs from larvae fed H₂O₂-supplemented food (Fig. 2B). These findings indicate that ROS positively regulates Akt phosphorylation.

**Figure 2.**
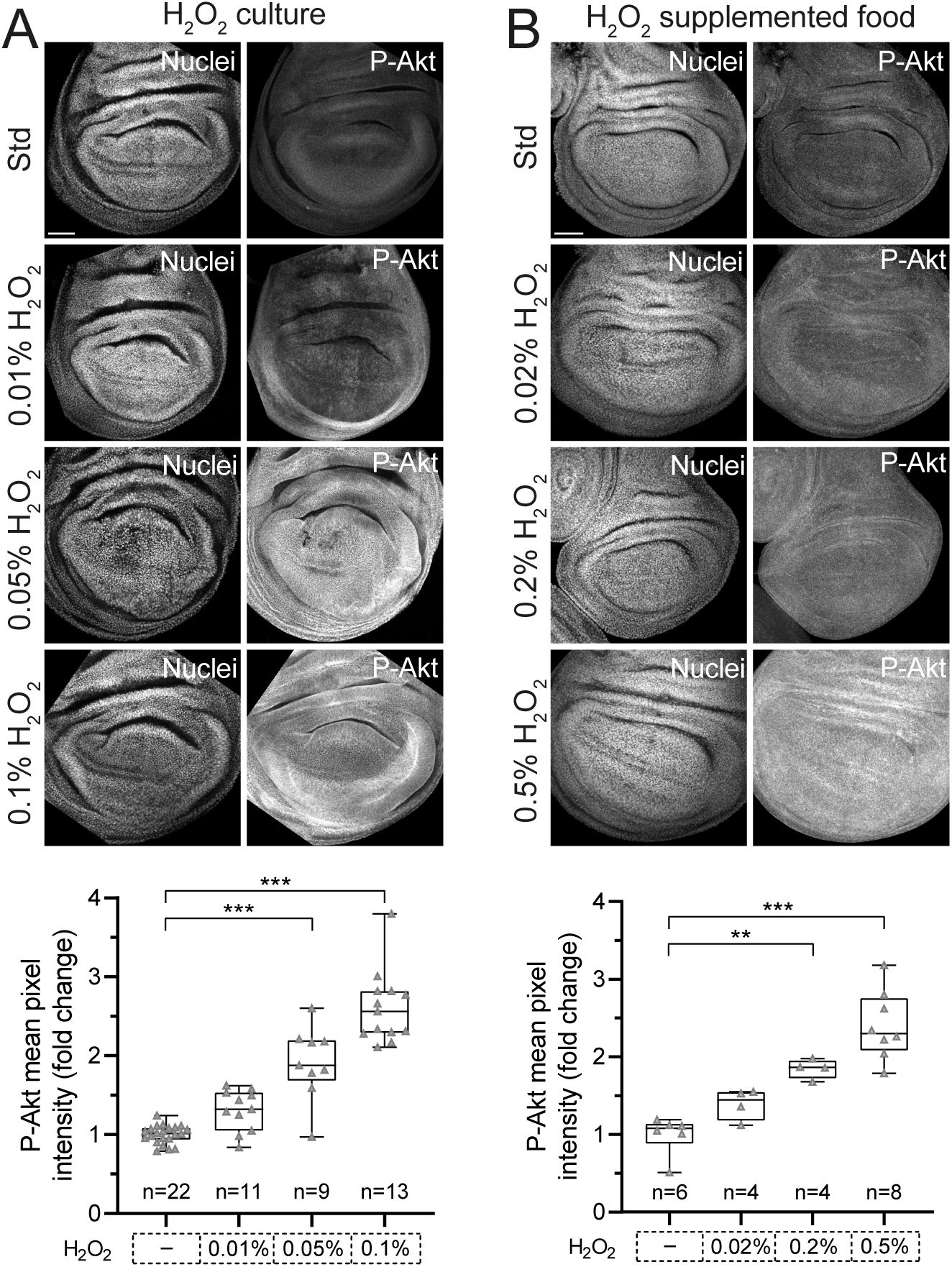
ROS induces phosphorylation of Akt in a dose-dependent manner. (A) Immunofluorescence detection of phospho-Akt in cultured wing imaginal discs after exposure to different concentrations of H_2_O_2_. (B) Immunofluorescence detection of phospho-Akt in wing imaginal discs after feeding larvae with standard food supplemented with different concentrations of H_2_O_2._ Quantification confirms that oxidative stress triggers Akt phosphorylation in a dose-dependent manner. Box plots show the maximum–minimum range (whiskers), the upper and lower quartiles (boxes), and the median value (horizontal line). Sample sizes are indicated in the graphs. Statistical significance was determined by one-way ANOVA followed by Tukey’s multiple-comparisons test: **p < 0.01 and ***p < 0.001. Images are representative of independent experiments. Scale bars 50µm.

### ROS inhibits PTEN in regenerating imaginal epithelia

These observations prompted us to examine whether the oxidative inhibition of PTEN regulates wing regeneration. To this end, we analyzed adult wings emerging from larvae that have been fed with or without H₂O₂ in different genetic backgrounds. Control animals were those in which genetic ablation, without any other genetic alteration, has been induced (*sal^E/Pv^>rpr; nub>+*). In these animals, wings from larvae raised on H₂O₂-supplemented food regenerated as in standard food, although they exhibited a modest reduction in size and a slight increase in the frequency of patterning defects (Fig. 3A, B and Supplementary Fig. 1). On the other hand, genetic ablation in the *sal* domain combined with overexpression of the wild type form of *Pten* in the *nubbin* domain *(sal^E/Pv^>rpr; nub>Pten^WT^)* led to a marked increase in aberrant wings with patterning defects, and a smaller size of the wings, indicating a regeneration failure. Remarkably, feeding animals of the same genotype with H₂O₂-supplemented food largely restored normal wing pattern and size (Fig. 3A, B). This recovery of the regenerative capacity was slightly stronger in female flies compared to males both in wing size and patterning. In females the normal regeneration dropped from 97.6 to 58% after *Pten* expression and recovered to 94% after feeding with H₂O₂-supplemented food. In males, the normal regeneration dropped from 92.9 to 55.6% after *Pten* expression and recovered to 83.7% after feeding with H₂O₂-supplemented food. These results indicate that ROS can counteract PTEN-mediated inhibition of regenerative growth, suggesting that redox regulation of PTEN is required for efficient tissue regeneration, an effect that is more pronounced in females.

**Figure 3.**
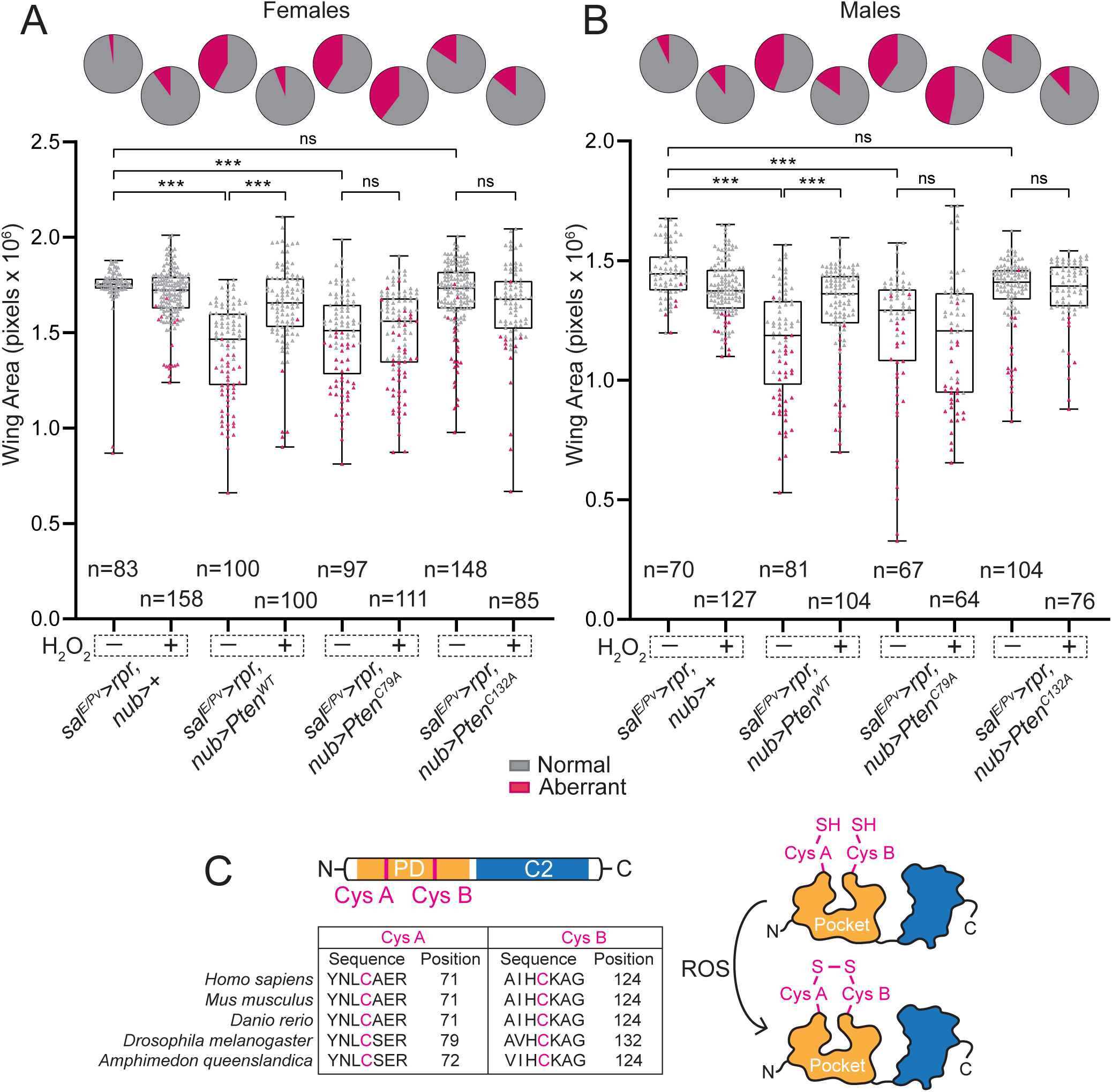
Oxidative stress targets Cys79 to inhibit PTEN activity. (A and B) Top: Pie charts showing the frequency of regenerated wings (gray) and non-regenerated or aberrant wings (purple). Bottom: Box plots showing adult wing area after exposure to H_2_O_2_. Adult female and male wings were analyzed separately. Box plots show the maximum–minimum range (whiskers), the upper and lower quartiles (boxes), and the median value (horizontal line). Genotypes and sample sizes are indicated below the plots. Statistical significance was determined by one-way ANOVA test followed by Tukey’s multiple-comparisons test was used: ns = not significant, ***p < 0.001. (C) Scheme of the PTEN protein indicating the phosphatase domain (PD), which contains the two Cys residues analyzed in this study. Sequence conservation and the positions of the corresponding Cys residues are indicated. PD: phosphatase catalytic domain. C2: membrane-binding domain that anchors PTEN to the membrane, positioning its catalytic site near membrane-associated substrates such as PIP_3_.

PTEN contains conserved cysteine residues that are susceptible to oxidative modification, which can reversibly impair its phosphatase activity (Kwon et al., 2004; Leslie et al., 2003). In *Drosophila*, Cys79 and Cys132 correspond to redox-sensitive cysteines identified in mammalian PTEN (Fig. 3C). Given the strong conservation of PTEN structure and function, these cysteines likely represent targets of ROS-mediated regulation in vivo. To investigate the contribution of these redox-sensitive cysteines to PTEN-dependent control of regenerative growth, we generated *UAS-*constructs carrying amino acid substitutions at the indicated cysteines (Fig. 3C). Throughout this study, these transgenes are referred to as *Pten* alleles.

To assess the functionality of these alleles, we analyzed Akt phosphorylation as a proxy for the phosphatase activity following expression of all transgenes under the control of *hh-Gal4* (*hh>*) driver, which restricts expression to the posterior compartment of the wing disc, allowing athe anterior compartment to serve as internal control. We found that phospho-Akt levels were reduced after overexpression of both the *Pten^C79A^* mutant and the *Pten^WT^* construct, whereas no reduction was observed following expression of *Pten^C132A^*, suggesting that the phosphatase activity has been compromised in the latter (Supplementary Figure 2).

Next, we tested whether these mutants affected the regeneration in a ROS-dependent manner. We did not observe remarkable regeneration defects after *Pten^C132A^* overexpression, either in standard conditions or in H₂O₂-supplemented food, which is consistent with the reduced catalytic activity of this form. In contrast, *UAS-Pten^C79A^* resulted in regeneration phenotypes comparable to those induced by *UAS-Pten^WT^* under standard feeding conditions, but no differences were detected in both female and male flies after H₂O₂-supplementation (Fig. 3A, B). This observation indicates that *UAS-Pten^C79A^*behaves as a phosphatase allele that is insensitive to changes in ROS levels. Based on these results, we focused all further experiments on *Pten^C79A^* as a variant that preserves phosphatase activity while being refractory to ROS-mediated regulation.

### PTEN oxidation at Cys79 promotes phosphorylation-dependent Akt activation in imaginal disc epithelia

Next, we examined how sensitivity to ROS affects Akt phosphorylation in *Pten^WT^* versus *Pten^C79A^* expressing cells in the posterior compartment (*hh>*). Wing discs were dissected and cultured in the presence or absence of H₂O₂ prior to antibody staining and quantification of confocal pixel intensity We found that in control discs expressing *UAS-GFP,* phospho-Akt levels were strongly increased throughout the tissue in response to H₂O₂ treatment. In *UAS-Pten^WT^* discs, oxidative stress induced Akt-phosphorylation, albeit in lower intensity than in the anterior compartment, consistent with ROS-mediated inhibition of PTEN activity. In contrast, in discs expressing *Pten^C79A^*, Akt phosphorylation was not induced by oxidative stress in the posterior compartment, whereas phospho-Akt levels showed an increase in the anterior control compartment (Fig. 4A). Together, these results indicate that ROS-mediated regulation of PTEN through Cys79 is required for Akt activation under oxidative conditions.

**Figure 4.**
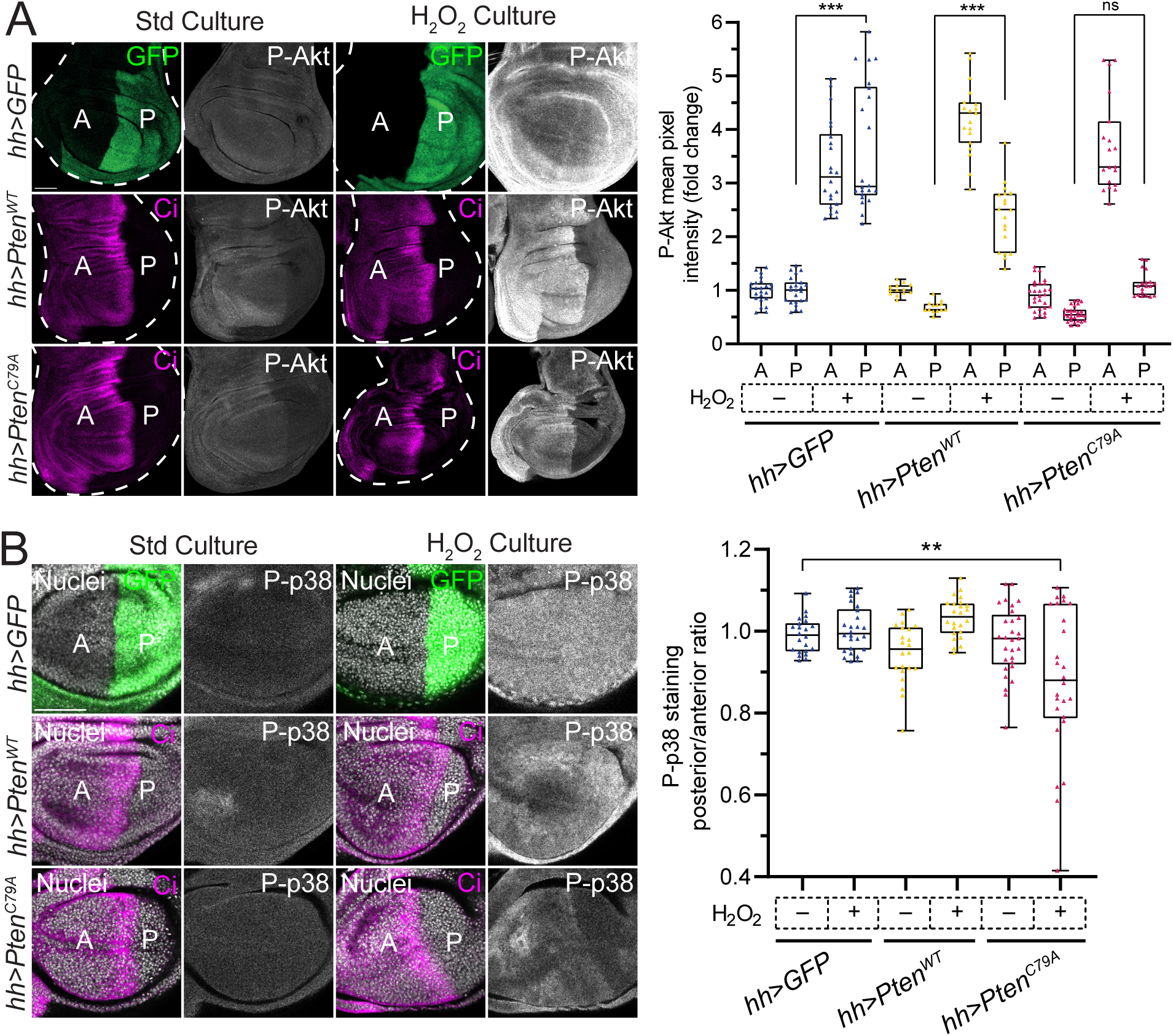
Oxidation of PTEN at Cys79 is required for ROS-dependent Akt and p38 activation in imaginal discs. (A) Representative images and quantification of phospho-Akt (P-Akt) levels in wing imaginal discs expressing GFP (control), *UAS-Pten^WT^*, or the ROS-insensitive *UAS-Pten^C79A^* allele in the posterior compartment (*hh>Gal4*), under standard culture conditions or following H₂O₂ treatment. Quantification shows the mean pixel intensity of P-Akt (fold change). Genotypes and sample sizes are as follows: *hh>GFP* std food (n=24), *hh>GFP* H_2_O_2_ food (n=22), *hh>Pten^WT^* std food (n=13), *hh>Pten^WT^* H_2_O_2_ food (n=19), *hh>Pten^C79A^* std food (n=26) and *hh>Pten^C79A^* H_2_O_2_ food (n=19). (B) Representative images and quantification of phospho-p38 (P-p38) levels under the same conditions. Compartments were identified by GFP fluorescence in control discs (*UAS-GFP*) or by immunostaining of anti-Ci, a marker of the anterior compartment, in discs expressing different transgenes (*UAS-Pten^WT^* and *UAS-Pten^C79A^*). Genotypes and sample sizes are as follows: *hh>GFP* std food (n=22), *hh>GFP* H_2_O_2_ food (n=26), *hh>Pten^WT^* std food (n=22), *hh>Pten^WT^*H_2_O_2_ food (n=24), *hh>Pten^C79A^* std food (n=29) and *hh>Pten^C79A^*H_2_O_2_ food (n=26). Box plots show the maximum–minimum range (whiskers), the upper and lower quartiles (boxes), and the median value (horizontal line). Statistical significance was determined by one-way ANOVA followed by Tukey’s multiple-comparisons test: ns = not significant, **p < 0.01 and ***p < 0.001. Images are representative of independent experiments. Scale bars 50µm.

Because ASK1-dependent activation of p38 requires Akt (Esteban-Collado et al., 2021, 2024; Santabárbara-Ruiz et al., 2019), we examined whether p38 phosphorylation was compromised after *Pten* manipulation. Given the variability in phospho-p38 staining between samples, we quantified p38 activation by calculating the ratio of posterior to anterior flourescence intensity for each disc. Overexpression of *Pten^WT^*or *Pten^C79A^* in the posterior compartment (*hh>*) in standard culture conditions did not result in detectable alterations in phospho-p38 relative to *UAS-GFP* controls. As previously described, culture of the imaginal discs in H₂O₂ resulted in strong p38 phosphorylation (Santabárbara-Ruiz et al., 2015). In discs expressing *Pten^WT^*, phospho-p38 levels were comparable between the anterior and posterior compartments. However, expression of *Pten^C79A^* resulted in significantly reduced p38 phosphorylation in the posterior compartment following H_2_O_2_ exposure, as evidenced by the decrease in the posterior to anterior P-p38 ratios (Fig. 4B). This observation is consistent with the ROS-insensitive phosphatase activity of the *Pten^C79A^*allele and suggests that PTEN Cys 79 oxidation is required for p38 activation under oxidative conditions.

### Stressors promote Akt and p38 activation through PTEN oxidation at Cys79 in the adult gut

The adult gut is an epithelium characterized by continuous cell turnover and a remarkable capacity to overcome damage and maintain tissue homeostasis (Micchelli & Perrimon, 2006; Ohlstein & Spradling, 2006). We therefore asked whether ROS-dependent Pten inactivation contributes to stress response in this tissue. To determine if Akt phosphorylation is responsive to oxidative stress in the adult intestine, we first analyzed Akt phosphorylation in flies fed with H₂O₂-supplemented food. Consistent with our findings in imaginal epithelia, phospho-Akt levels increased in adult guts in a dose-dependent manner following H₂O₂ treatment (Fig. 5A).

**Figure 5.**
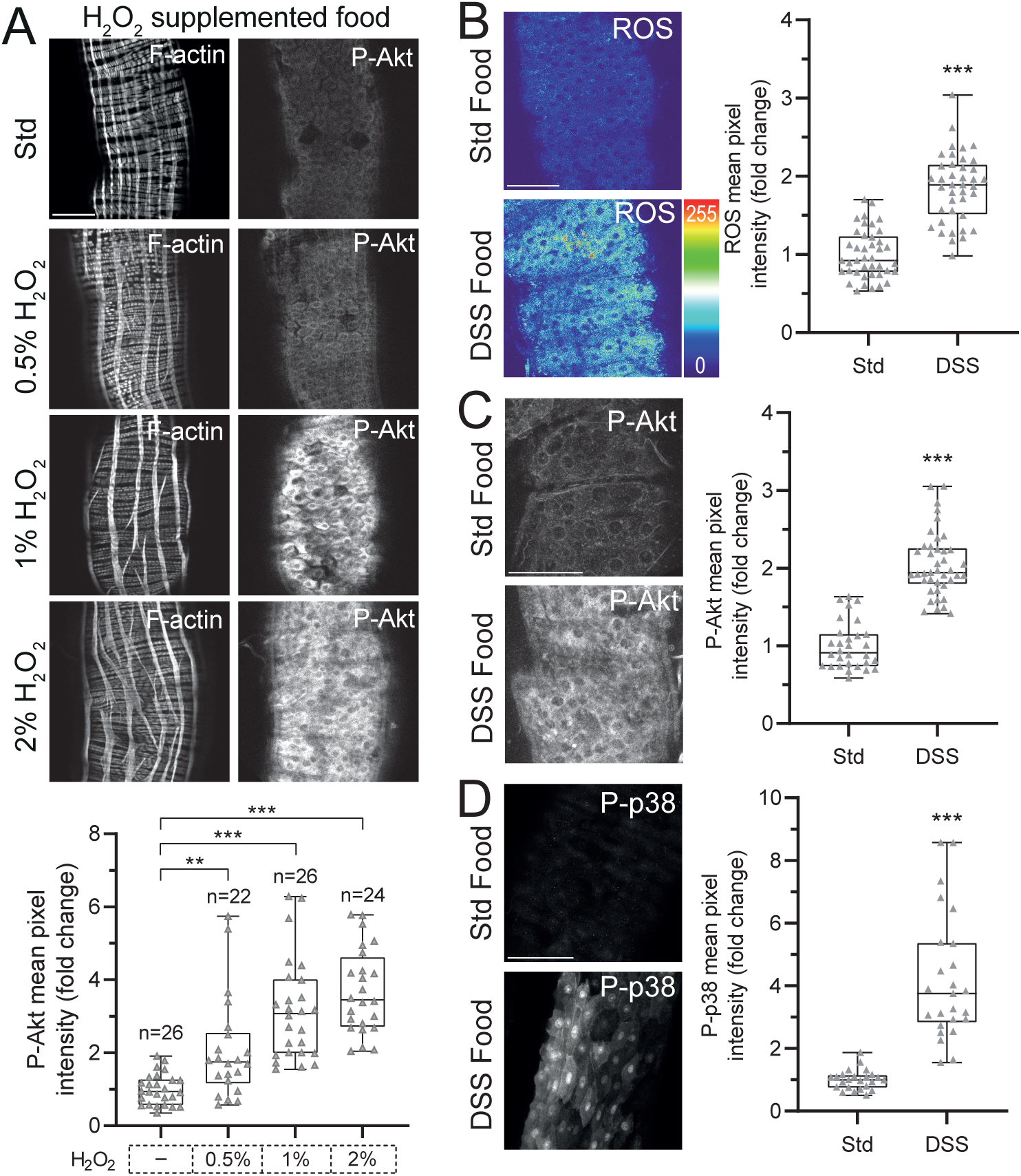
Damage-induced ROS production correlates with Akt and p38 activation in the adult midgut. (A) Phospho-Akt staining in adult midguts from flies fed standard food supplemented with increasing concentrations of H_2_O_2_. Quantification of P-Akt (mean pixel intensity, fold change) confirms that ROS induces Akt phosphorylation in a dose-dependent manner. Sample sizes are indicated in the graph. Statistical significance was determined by one-way ANOVA followed by Tukey’s multiple-comparisons test: **p < 0.01 and ***p < 0.001. Images are representative of independent experiments. (B) Representative images of ROS levels in adult midguts under standard conditions or following DSS-induced damage. Quantification of ROS signal (mean pixel intensity, fold change) confirms increased oxidative stress after tissue damage. Sample sizes: std food (n=41) and DSS food (n=40). (C) Representative images of phosphorylated Akt (P-Akt) in adult midguts under standard conditions or following DSS-induced damage. Quantification of P-Akt levels (mean pixel intensity, fold change) shows increased Akt phosphorylation upon damage. Sample sizes: std food (n=30) and DSS food (n=41). (D) Representative images of phosphorylated p38 (P-p38) in adult midguts under standard conditions or following DSS-induced damage. Quantification of P-p38 levels (mean pixel intensity, fold change) shows increased p38 phosphorylation upon damage. Sample sizes: std food (n=24) and DSS food (n=25). Box plots show the maximum–minimum range (whiskers), the upper and lower quartiles (boxes), and the median value (horizontal line). Two-tailed Student’s t-test was used for statistical comparisons (B, C, D): ***p < 0.001. Images are representative of independent experiments. Scale bars 50µm.

Among the injury paradigms commonly used in the gut, dextran sodium sulfate (DSS) induces inflammation resembling ulcerative colitis in mammals (Kawada, 2007) and causes epithelial damage in *Drosophila* leading to intestinal barrier disruption, increased intestinal stem cell (ISC) division and enteroblasts production and sometimes lethality (Amcheslavsky et al., 2009). Consistent with the induction of oxidative stress, DSS treatment led to robust ROS production and was accompanied by strong activation of both Akt and p38 in damaged adult intestines (Fig. 5B-D). These observations indicate that tissue damage is associated with activation of growth signaling pathways in the adult gut, prompting us to investigate whether ROS-dependent inhibition of PTEN underlies Akt activation during the intestinal damage response.

We next examined how oxidative stress affects the proliferative response of ISCs expressing the different *Pten* alleles, as modulation of the redox balance in the intestinal epithelium is known to strongly influence ISC proliferation. (Hochmuth et al., 2011). We induced the oxidative stress by feeding the animals with H₂O₂-supplemented food, and ISCs and enteroblasts were visualized using the *esg-Gal4, UAS-GFP* reporter (*esg>GFP*). We observed a marked increase in the number of *esg>GFP*-positive cells (Fig. 6A-B) and mitoses (Fig. 6C) after H₂O₂ intake. Expression of *Pten^WT^*under the *esg-Gal4* driver rsulted in a significant reduction of mitoses (Fig. 6C) and a weak decrease of *esg>GFP*-positive cells under standard feeding conditions. Upon H₂O₂ supplementation, guts expressing *Pten^WT^* exhibited an increased number of *esg>GFP*-positive cells and mitoses (Fig. 6A-C), although to a lesser extent than observed in controls. In contrast, expression of the ROS-insensitive *Pten^C79A^* allele, which also reduced the number of mitoses and GFP-positive cells under control conditions, largely abolished the proliferative response to H₂O₂ exposure (Fig. 6A-C). These observations indicate that *Pten^C79A^* mutant is refractory to ROS-mediated regulation and link ISC proliferative responses to oxidative stress with PTEN/Akt signaling.

**Figure 6.**
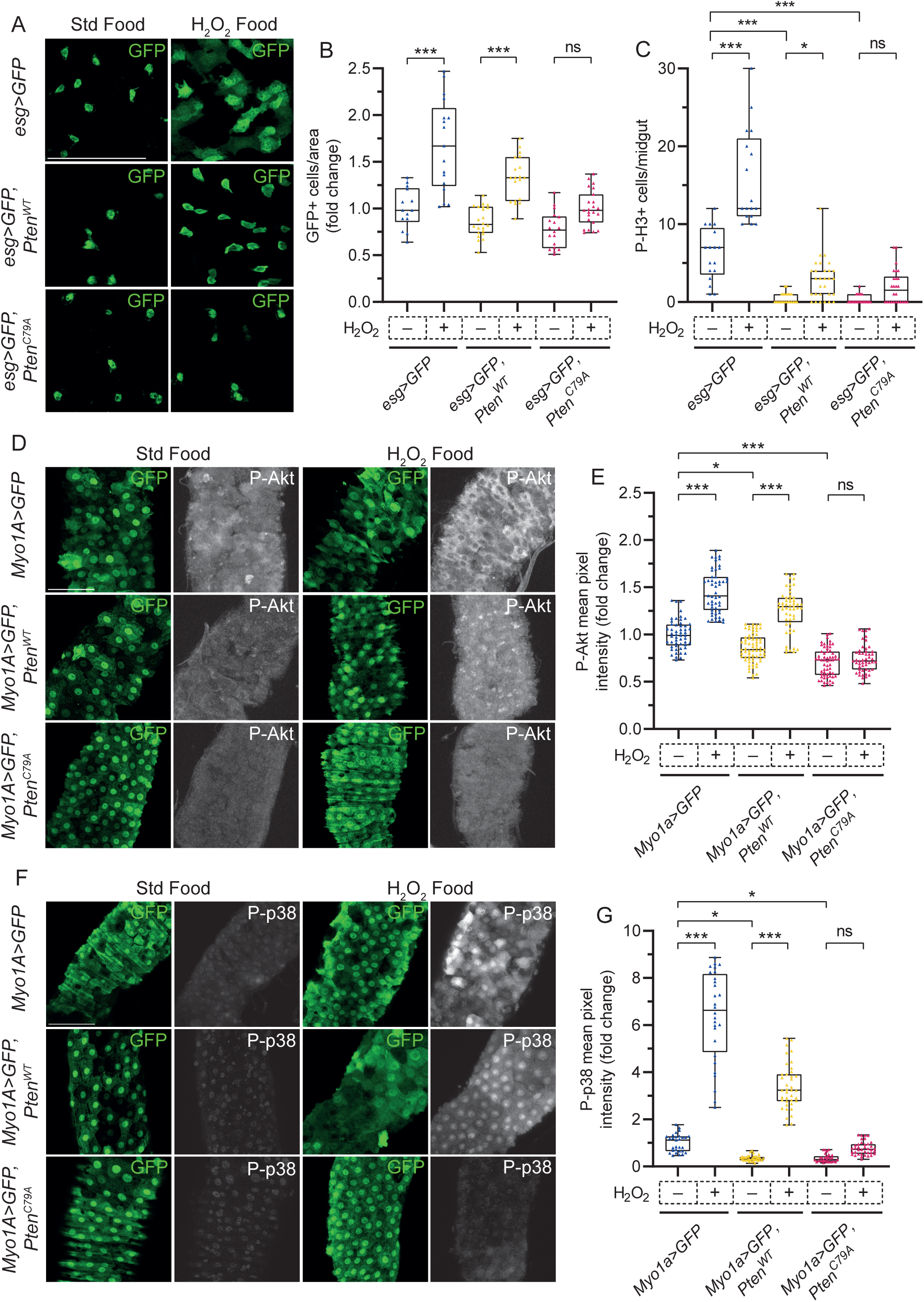
ROS regulates ISC proliferation and Akt/p38 signaling through PTEN inactivation in the adult midgut. (A, B) Representative images (A) and quantification (B) of *esg>GFP*-positive intestinal stem cells (ISCs) and enteroblasts in adult midguts under standard food or H₂O₂- supplemented food for the indicated genotypes. Genotypes and sample sizes are as follows: *esg>GFP* std food (n=14), *esg>GFP* H_2_O_2_ food (n=17), *esg>GFP,Pten^WT^* std food (n=22), *esg>GFP,Pten^WT^*H_2_O_2_ food (n=19), *esg>GFP,Pten^C79A^* std food (n=28) and *esg>GFP,Pten^C79A^* H_2_O_2_ food (n=23). (C) Quantification of mitotic cells by phospho-histone H3 (P-H3) under the same conditions. Genotypes and sample sizes are as follows: *esg>GFP* std food (n=17), *esg>GFP* H_2_O_2_ food (n=17), *esg>GFP,Pten^WT^* std food (n=23), *esg>GFP,Pten^WT^*H_2_O_2_ food (n=29), *esg>GFP,Pten^C79A^* std food (n=23) and *esg>GFP,Pten^C79A^*H_2_O_2_ food (n=26). (D) Representative images of phospho-Akt (P-Akt) staining in enterocytes (*Myo1A>GFP*) under standard food or H₂O₂- -supplemented food conditions. (E) Quantification of P-Akt levels in enterocytes expressing *Pten^WT^* or *Pten^C79A^*. Genotypes and sample sizes are as follows: *Myo1A>GFP* std food (n=53), *Myo1A>GFP* H_2_O_2_ –supplemented food (n=51), *Myo1A>GFP,Pten^WT^* std food (n=67), *Myo1A>GFP,Pten^WT^*H_2_O_2_ –supplemented food (n=52), *Myo1A>GFP,Pten^C79A^*std food (n=60) and *Myo1A>GFP,Pten^C79A^* H_2_O_2_ –supplemented food (n=50). (F) Representative images of phospho-p38 (P-p38) staining in enterocytes under the indicated conditions. (G) Quantification of P-p38 levels in enterocytes expressing *Pten^WT^* and *Pten^C79A^*. Genotypes and sample sizes: *Myo1A>GFP* std food (n=29), *Myo1A>GFP* H_2_O_2_ –supplemented food (n=28), *Myo1A>GFP,Pten^WT^* std food (n=36), *Myo1A>GFP,Pten^WT^* H_2_O_2_ –supplemented food (n=39), *Myo1A>GFP,Pten^C79A^* std food (n=30) and *Myo1A>GFP,Pten^C79A^* H_2_O_2_ –supplemented food (n=38). Box plots show the maximum–minimum range (whiskers), the upper and lower quartiles (boxes), and the median value (horizontal line). Statistical significance was determined by one-way ANOVA followed by Tukey’s multiple-comparisons test: ns = not significant, *p < 0.05 and ***p < 0.001. Images are representative of independent experiments. Scale bars 50µm.

We then investigated whether ROS–PTEN signaling also regulates Akt activity in the adult gut. To this end, we focused on differentiated enterocytes, rather than the ISCs population, using the enterocyte-specific driver *Myo1A-Gal4*. Enterocytes constitute the majority of the adult midgut epithelium and arise from ISC-derived enteroblasts that terminally differentiate into polyploid absorptive cells expressing Myosin31DF (Myo1A) (Hung et al., 2021; Neophytou et al., 2025). Because enterocytes are highly abundant, we reasoned that the *Myo1A-Gal4* driver would be more sensitive than the *esg-Gal4* driver for detecting tissue-wide signaling changes. Following H₂O₂ feeding, phospho-Akt levels were strongly increased throughout the midgut epithelium (Fig. 6D-E). Expression of *Pten^WT^* in enterocytes attenuated phospho-Akt levels under standard conditions, whereas simultaneous H₂O₂ exposure increased phospho-Akt, consistent with ROS-mediated inhibition of PTEN activity (Fig. 6D-E). In contrast, expression of *Pten^C79A^* abolished H₂O₂-induced Akt phosphorylation in enterocytes (Fig. 6D-E). Together, these results indicate that ROS regulate Akt signaling in the adult gut by modulating PTEN activity.

Finally, we examined whether p38 activation is affected by PTEN manipulation. Oxidative stress induced by H₂O₂ feeding in wild-type flies triggered robust p38 phosphorylation, as reported (Patel et al., 2019). Expression of *Pten^WT^* or *Pten^C79A^*in enterocytes reduced basal phospho-p38 levels (Fig. 6F). Upon H₂O₂ exposure, *Pten^WT^*-expressing guts showed increased p38 phosphorylation, whereas *Pten^C79A^-*expressing guts failed to respond to oxidative stress. These results indicate that ROS-dependent activation of p38, required for regeneration, is regulated by PTEN.

## DISCUSSION

In this study, we identify redox regulation of PTEN as a conserved mechanism linking oxidative stress to Akt activation during tissue regeneration and homeostasis in *Drosophila*. We show that ROS levels positively correlate with Akt phosphorylation in both regenerating imaginal discs and the adult midgut, and that this response depends on PTEN activity. Using targeted expression of a redox-insensitive *Pten* allele, we demonstrate that Cys79 is required for ROS-mediated inhibition of PTEN and subsequent activation of Akt signaling. These findings provide in vivo evidence that redox regulation of PTEN directly controls growth and regenerative responses across distinct tissues and developmental stages.

Functionally, our results indicate that ROS-mediated modulation of PTEN activity is required for coordinating cellular responses to tissue damage. In imaginal discs, this mechanism promotes regenerative growth following apoptosis, while in the adult midgut it supports both ICS proliferation and Akt activation in differentiated enterocytes. The inability of the *Pten^C79A^* allele to respond to oxidative stress uncouples Akt activation from ROS levels, resulting in impaired regenerative and proliferative responses despite preserved phosphatase activity. Together, these data support a model in which transient ROS production acts as a physiological signal that locally inhibits PTEN, thereby enabling Akt-dependent growth and repair. Given the high degree of conservation of PTEN structure and its redox sensitivity, this mechanism is likely to represent a broadly used strategy by which tissues integrate metabolic and damage-induced signals to regulate regeneration and maintain homeostasis (Trinh et al., 2024).

The uncoupling of the oxidative sensitivity from phosphatase activity in *Pten^C79A^* contrasts with *Pten^C132A^*. The reduced catalytic activity of the *Pten^C132A^* is consistent with previous studies showing that substitution of the equivalent Cys^124^ residue to human PTEN abolishes phosphatase activity and promotes tumorigenic growth (Chao et al., 2020; Myers et al., 1997, 1998; Post et al., 2020). Mutation in the Cys^124^ residue generates a catalytic inactive form of PTEN that results in an accumulation of PIP_3_ (Myers et al., 1998).

Redox regulation of PTEN has been extensively documented in mammalian cells, where transient oxidation of conserved cysteine residues promotes PI3K/Akt signaling during growth factor stimulation, wound repair, and inflammatory responses (Burge et al., 2025; Kwon et al., 2004; Leslie et al., 2003; Trinh et al., 2024). Our in vivo data in *Drosophila* support the idea that this regulatory mechanism is evolutionarily conserved and operates at the level of intact tissues to coordinate tissue regeneration and homeostasis. Notably, dysregulated ROS–PTEN–PI3K/Akt axis has been implicated in a wide range of pathological conditions such as cancer, metabolic disorders, and age-associated tissue decline in mammals (Cheung & Vousden, 2022; Ranbhise et al., 2025; Sies & Jones, 2020; Zhang et al., 2020).

The role of PI3K/Akt signaling in regeneration has been extensively documented across vertebrate tissues, where it contributes to processes such as tendon repair, axonal regeneration, and liver regeneration (Goto et al., 2025; Jung et al., 2021; Wang et al., 2022). In addition, ROS have been implicated in PI3K/Akt-dependent regenerative responses, including mammalian axonal regeneration (Hervera et al., 2018). Injury-induced ROS production is a conserved feature of tissue damage, and H₂O₂ has been shown to regulate MAPK signaling during wound healing across multiple systems (Hunt et al., 2024). Our findings that damage-induced ROS inactivate PTEN, thereby relieving its inhibition of Akt and promoting p38 activation, suggest that this mechanism may represent a conserved strategy for coordinating regenerative responses beyond *Drosophila*.

It is generally assumed that ROS production following tissue damage is transient, as tissue repair limits further ROS generation (Dunnill et al., 2017). In addition, oxidation of PTEN is reversible, allowing its phosphatase activity to be restored once oxidative stress declines (Kwon et al., 2004; Lee et al., 2002). Together, these features suggest a mechanism in which transient ROS production induces sufficient Akt activation to trigger a repair response, including activation of stress-responsive kinases such as p38 or JNK. The transient and reversible nature of this response enables these kinases to promote tissue repair while minimizing detrimental consequences associated with sustained or excessive signaling. This provides a framework in which redox-dependent modulation of PTEN couples transient oxidative stress to the controlled activation of regenerative signaling pathways.

In this context, our findings, together with previous work, support a model in which ROS coordinate multiple regulatory nodes within the stress response (Fig. 7). First, ROS promote the dissociation of thioredoxin from ASK1, enabling its activation (Patel et al., 2019; Santabárbara-Ruiz et al., 2019; Shiizaki et al., 2013; Takeda et al., 2008). Second, ROS act on ligand-independent TNFR signaling to facilitate TRAF recruitment to ASK1, thereby promoting downstream p38 activation (Esteban-Collado et al., 2024). Third, as shown here, ROS-mediated inhibition of PTEN, enhances Akt phosphorylation, which in turn modulates ASK1 activity to bias signaling towards p38-dependent survival and repair responses (Esteban-Collado et al., 2021).

**Figure 7.**
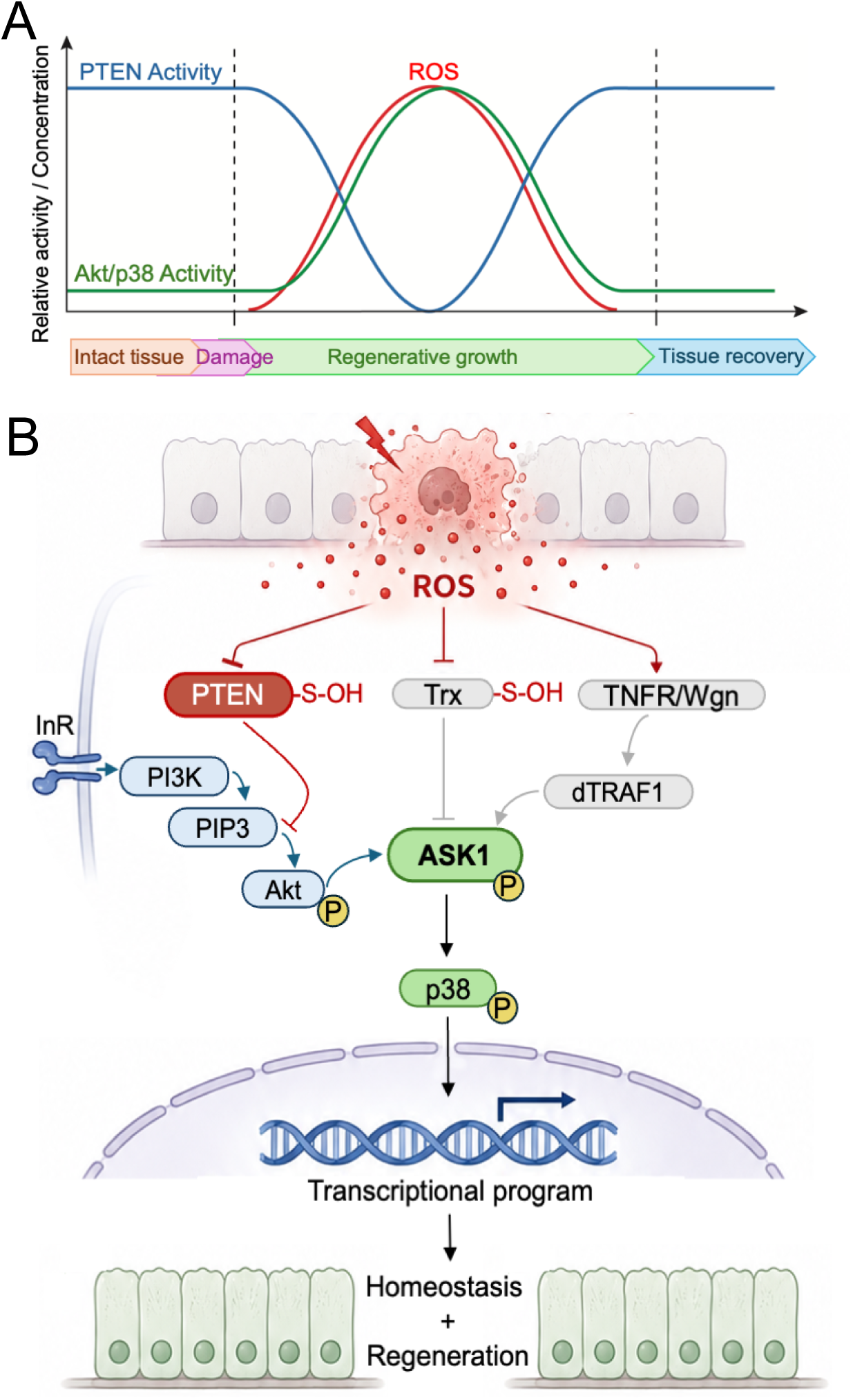
ROS coordinate PTEN/PI3K/Akt and ASK1 signaling to control regenerative responses. (A) Model illustrating the temporal sequence of events during the damage response. Transient ROS production following tissue damage (red) generates an oxidative environment that reversibly inhibits PTEN (blue). This results in a transient increase in Akt and p38 activity (green), thereby triggering a repair response (regenerative growth). Shortly after oxidation, PTEN activity is recovered. This returns Akt and p38 activity to basal levels, preventing the detrimental effects of sustained activation. This mechanism allows for precise spatiotemporal fine tuning of the damage response and the subsequent recovery of tissue homeostasis. (B) Model illustrating how reactive oxygen species (ROS) coordinate multiple signaling pathways to regulate tissue repair. ROS act on three key regulatory nodes. First, ROS promote oxidation and inhibition of PTEN, relieving its suppression of PI3K signaling and enabling Akt activation. Second, ROS induce dissociation of thioredoxin (Trx) from ASK1, facilitating ASK1 activation. Third, ROS enhance ligand-independent TNFR signaling, promoting TRAF-dependent activation of ASK1 required for p38 activation. Activated Akt modulates ASK1 signaling output, biasing downstream pathway selection towards p38-dependent survival and regenerative responses. Together, these interconnected mechanisms position ROS as a central integrator of stress signaling, coordinating the activation of regenerative pathways while ensuring that signaling remains tightly controlled.

Together, these mechanisms position ROS as a central integrator of stress signaling, coordinating ASK1 activation and downstream pathway selection to ensure an effective yet controlled regenerative response while limiting the detrimental consequences of excessive or sustained signaling.

The demonstration that redox-sensitive inhibition of PTEN is required for appropriate regenerative responses in ISC, enterocytes and imaginal discs raises the possibility that similar mechanisms underlie tissue repair and adaptive growth in more complex organisms. Thus, *Drosophila* provides a genetically tractable model to dissect how physiological and pathological ROS levels are interpreted through PTEN, offering insights into the mechanisms by which redox signaling regulates tissue regeneration and disease progression, which may guide the development of therapeutic strategies targeting these pathways in mammalian tissues.

## Acknowledgements

The authors would like to thank Dr. Manel Bosch from the Optical Microscopy Unit of the CCiT of the Universitat de Barcelona for his assistance. This research was funded by PID2024-158952NB- I00 from MICIU/AEI/10.13039/501100011033/FEDER, UE and by the Agència de Gestió d’Ajuts Universitaris i de Recerca of the Generalitat de Catalunya (2021SGR00293) to FS and MC.

## MATERIALS AND METHODS

### Drosophila Strains

Animals were reared on standard fly food. The *Drosophila melanogaster* strains *sal^E/Pv^-LHG* and *lexO-rpr* have been previously described (Santabárbara et al. 2015). Other strains used in this study were: *UAS-Pdk1,Akt1* (Rintelen et al., 2001), *esg-Gal4*, M*yo1A-Gal4*, *tub-Gal80^TS^*and *UAS-GFP* (gift from M. Milán, IRB Barcelona) and *nub-Gal4* and *hh-Gal4* (described in Flybase). The strains *UAS-Pten^WT^*, *UAS-Pten^C79A^* and *UAS-Pten^C132A^*were generated as described below. The following strains were obtained from Bloomington Drosophila Stock Center (BDSC): *tub-Gal80^TS^*(RRID: BDSC_7017) and *UAS-GFP* (RRID: BDSC_4776). The *w^1118^;+;+* strain was used as a control strain. A complete list of genotypes is provided below.

### Genotypes and Nomenclature

Abbreviated genotype names used throughout the manuscript are listed below along with their corresponding full genotypes.

Fig. 1 (B,C)

*sal^E/Pv^>rpr, nub>+*: *w; nub-Gal4:lexO-rpr/+; sal^E/Pv^-lexA:tub-Gal80^TS^/+*

*sal^E/Pv^>rpr, nub>Pdk1,Akt1*: *w; nub-Gal4:lexO-rpr/GMR-Gal4:UAS-Pdk1:UAS-Akt1; sal^E/Pv^-lexA:tub-Gal80^TS^/+*

Fig. 2 (A,B)

*w^1118^; +; +*

Fig. 3 (A,B)

*sal^E/Pv^>rpr, nub>+*: *w; nub-Gal4:lexO-rpr/+; sal^E/Pv^-lexA:tub-Gal80^TS^/+*

*sal^E/Pv^>rpr, nub>Pten^WT^*: *w; nub-Gal4:lexO-rpr/UAS-Pten^WT^; sal^E/Pv^-lexA:tub-Gal80^TS^/+*

*sal^E/Pv^>rpr, nub>Pten^C79A^*: *w; nub-Gal4:lexO-rpr/UAS-Pten^C79A^; sal^E/Pv^-lexA:tub-Gal80^TS^/+*

*sal^E/Pv^>rpr, nub>Pten^C132A^*: *w; nub-Gal4:lexO-rpr/UAS-Pten^C132A^; sal^E/Pv^-lexA:tub-Gal80^TS^/+*

Fig. 4 (A,B)

*<u>hh>GFP</u> stands for w; tub-Gal80^TS^/UAS-GFP; hh-Gal4/UAS-GFP*

*hh>Pten^WT^*: *w; tub-Gal80^TS^/UAS-Pten^WT^; hh-Gal4/+*

*hh>Pten^C79A^*: *w; tub-Gal80^TS^/UAS-Pten^C79A^; hh-Gal4/+*

Fig. 5 (A,B,C,D)

*w^1118^; +; +*

Fig. 6 (A,B,C)

*esg>GFP*: *w; esg-Gal4:UAS-GFP/+; tub-Gal80^TS^/+*

*esg>Pten^WT^*: *w; esg-Gal4:UAS-GFP/UAS-Pten^WT^; tub-Gal80^TS^/+*

*esg>Pten^C79A^*: *w; esg-Gal4:UAS-GFP/UAS-Pten^C79A^; tub-Gal80^TS^/+*

Fig. 6 (D,E,F,G)

*Myo1A>GFP*: *w; Myo1A-Gal4/+; tub-Gal80^TS^:UAS-GFP /+*

*Myo1A>Pten^WT^*: *w; Myo1A-Gal4/UAS-Pten^WT^; tub-Gal80^TS^:UAS-GFP /+*

*Myo1A>Pten^C79A^*: *w; Myo1A-Gal4/UAS-Pten^C79A^; tub-Gal80^TS^:UAS-GFP /+*

Suppl. Fig. 2 (A,B)

*<u>hh>GFP</u>: w; tub-Gal80^TS^/UAS-GFP; hh-Gal4/UAS-GFP*

*hh>Pten^WT^*: *w; tub-Gal80^TS^/UAS-Pten^WT^; hh-Gal4/+*

*hh>Pten^C79A^*: *w; tub-Gal80^TS^/UAS-Pten^C79A^; hh-Gal4/+*

*hh>Pten^C132A^*: *w; tub-Gal80^TS^/UAS-Pten^C132A^; hh-Gal4/+*

### Genetic ablation and dual Gal4/lexA transactivation system for wing regeneration analysis

Adult wing regeneration was analyzed using the combined action of *Gal4/UAS* and *lexA/LexO* binary systems as previously described (Esteban-Collado et al., 2021; Santabárbara-Ruiz et al., 2019). Briefly, the lexA activation domain was replaced by the Gal4 activation domain to enable repression by the Gal80, a well-known inhibitor of Gal4 activity. This modified lexA protein (LexA-Hinge-Gal4 activation domain, hereafter LHG) (Yagi et al., 2010) was used to achieve spatiotemporal control of the expression of the pro-apoptotic gene *reaper* (*lexO-rpr*) under the control of the *sal^E/Pv^* regulatory region, which is specifically active in the central zone of the wing imaginal disc (genetic ablation). In addition, the *Gal4/UAS* system was used for simultaneous expression of the desired transgenes throughout the entire wing pouch with the *nub-Gal4* driver. The temperature-sensitive allele *tubGal80^TS^* allowed temporal control of both expression systems within the wing disc without affecting the rest of the organism.

Genetic ablation was induced during larval stages using the genotype *sal^E/Pv^-LHG, lexO-rpr, tubGal80^TS^*. Adult flies were allowed to lay eggs for 6 h at 17°C. The offspring were maintained at 17°C until day 8 of development (192 h after egg laying), when temperature was switched to 29°C for 11h to induce genetic ablation. After that, they were returned to 17°C until adults emerged. Adult flies were collected and fixed in glycerol:ethanol (1:2 mix) for 24 h. Wings were dissected in distilled water and washed in 100% ethanol before mounting in lactic acid:ethanol (6:5 mix). After mounting, gentle pressure was applied on top the coverslip for 24 h, and preparations containing wing preparations were photographed using a Leica DMLB microscope.

Wing areas were measured using FIJI/ImageJ. Wing phenotypes were classified by visual inspection into two categories. The normal regeneration category included wings displaying a normal set of veins and interveins, as well as wings lacking only a segment of the anterior vein L2. The aberrant regeneration category included wings with partially or completely missing veins and/or interveins (Supplementary Figure 1). Female and male wings were analyzed independently.

### Nutrient restriction conditions

Standard food contains fresh yeast (64 g/L), dextrose (64 g/L), wheat flour (40 g/L), Bacto agar (8.8 g/L). Nutrient restricted food (or 10% yeast food) was prepared by reducing the amount of yeast to 6.4 g/L without altering the other ingredients. All crosses and experiments were performed under non-crowding conditions.

Nutrient restriction experiments were performed as follows. Embryos were maintained at 17°C. On day 7 of development (168 h after egg laying), larvae were removed from standard food, washed in PBS and transferred to vials containing 10% yeast. Control animals were subjected to the same handling procedure but were transferred to fresh standard food instead. On day 8 (192 h after egg laying), larvae were transferred to 29°C for 11 h to activate transgene expression, as the Gal80 inhibitor is inactive at this temperature. Subsequently, larvae were returned to 17°C until they reached adulthood. Thus, nutrient restriction was imposed starting 24 h before cell death induction, throughout the cell ablation period and during the entire regeneration process until adulthood. To exclude any toxicity due to the insertion of the transgene, all experiments were carried out in parallel under constant 17 °C conditions, to maintain *tubGal80^TS^* activity and prevent transgene (*UAS*-or *lexO*-mediated) expression. Under these conditions, no wing patterning defects were detected.

### Oxidative stress induction

Three different assays were used to induce oxidative stress:

1. ROS exposure to cultured imaginal discs. Wing discs were dissected in Schneider’s *Drosophila* Medium (VWR Chemicals Cat.# 392-0419), transferred to Schneider’s medium containing H_2_O_2_ (0.01%, 0.05% and 0.1% for Figure 2; 0.1% for Figure 4) for 15 min, washed in Schneider’s medium and fixed in 4% paraformaldehyde (Electron Microscopy Sciences Cat.# 15710) in PBS). Samples were processed using standard immunostaining methods.
2. ROS-supplemented food for imaginal disc analysis. Third-instar larvae maintained at 17°C on standard food were transferred on day 7 (168 h after egg laying) to vials containing H_2_O_2_ in the fly food (0.02%, 0.2% and 0.5% for Figure 2; 0.02% for Figure 3). Control larvae were transferred to vials with standard food. On day 8 (192 h after egg laying), imaginal discs were dissected and processed using standard immunostaining methods. For wing regeneration experiments, larvae were shifted to 29°C for 11 h on day 8 to induce transgene expression and then transferred to 17°C until adulthood (as described above).
3. ROS-supplemented food for the gut analysis. Young, mated females (from 4 to 7 days old) were raised at 17°C and transferred to vials with H_2_O_2-_supplemented food (0.5%, 1% and 2% for Figure 5; 2% for Figure 6). Flies were maintained at 29°C to induce transgene expression for 48h before dissection. Control flies were transferred to vials containing standard food. Flies were transferred to fresh H_2_O_2_ supplemented food every 24 h. Adult midguts of were dissected and processed for analysis.

To avoid loss of oxidative capacity, H_2_O_2_ (Sigma-Aldrich Cat.# 516813) was added to the food at a temperature below 40°C. H_2_O_2_ supplemented food was freshly made before use.

### DSS exposure

Young, mated females (from 4 to 7 days old) were raised at 17°C and transferred to food supplemented with 3% DSS or standard food (control) at 29°C for 48 h before dissection. Flies were transferred every 24 h to vials with new DSS food. DSS (Sigma-Aldrich Cat.# 42867) was added to the food at a temperature below 40°C. DSS supplemented food was freshly prepared before use.

### ROS detection by CellROX staining

Adult midguts were dissected in Schneider’s medium, incubated for 30 min in 5 μM CellROX™ Deep Red Reagent (Thermo Fisher Scientific, Cat. #C10422), fixed for 30 min in 4% PFA, washed in 1× PBS, and mounted in Vectashield Plus Antifade Mounting Medium (Vector Laboratories, Cat. #H-1900). Imaging was performed within 1 h of mounting. Control (without DSS) and experimental (with DSS) samples were processed and imaged in parallel.

### Immunofluorescence

The primary antibodies used were against P-Akt S473 (rabbit 1:200; Cell Signaling Technology Cat.# 4060, RRID: AB_2315049), P-p38 T180/Y182 (rabbit 1:50; Cell Signaling Technology Cat.# 9211, RRID: AB_331641), P-H3 S10 (mouse 1:400; Cell Signaling Technology Cat.# 9706, RRID: AB_331748), cleaved *Drosophila* Death Caspase-1 (Dcp-1) D215 (rabbit 1:200; Cell Signalling Technology Cat.# 9578, RRID: AB_2721060), Ci (rat 1:100; DSHB (Iowa) Cat.# 2A1, RRID: AB_2109711), and GFP (chicken 1:1000; Abcam Cat.# AB13970, RRID: AB_300798). The fluorescently labeled secondary antibodies were used 1:200 in PBS 0.3% Triton X-100 (PBT 0.3%) for wing imaginal discs and in PBS 0.1% Triton X-100 (PBT 0.1%) for adult midguts. The following secondary antibodies were used: donkey anti-rabbit Alexa Fluor 568 (Thermo Fisher Scientific Cat.# A10042, RRID: AB_2534017), goat anti-rat Alexa Fluor 488 (Thermo Fisher Scientific Cat.# A11006, RRID: AB_2534074), goat anti-mouse Alexa Fluor 568 (Thermo Fisher Scientific Cat.# A11004, RRID: AB_2534072) and goat anti-chicken Alexa Fluor 488 (Thermo Fisher Scientific Cat.# A11039, RRID: AB_2534096).

Wing imaginal discs from late third-instar larvae were dissected in Schneider’s medium, fixed in 4% paraformaldehyde (PFA) for 50min at 4°C, washed in PBT 0.3%, blocked for 30min in PBT 0.3% containing 2% BSA (Sigma-Aldrich Cat.# A9647) and incubated overnight with primary antibody at 4°C. The following day, discs were washed in 0.3% PBT and incubated with secondary antibodies for 2h at room temperature. After washing, discs were stained for 30min at room temperature with nuclear markers DAPI (5μg/mL; Thermo Fisher Scientific Cat.# D21490) or TOPRO-3 (1μM; Thermo Fisher Scientific Cat.# T3605). Discs were mounted in SlowFade Diamond Antifade Mountant (Thermo Fisher Scientific Cat.# S36967).

Adult midguts were dissected in cold filtered 1x PBS (pH 7.4), fixed in 4% PFA for 1h at 4°C, washed in 0.1% PBT, blocked for 1h in 0.1% PBT containing 4% BSA and incubated overnight with primary antibody at 4°C. The following day, midguts were washed in PBT 0.1% and incubated with secondary antibody for 1h at room temperature. After washing, midguts were stained for 30min at room temperature with Phalloidin Alexa Fluor 488 (1:400, Thermo Fisher Scientific Cat.# A12379) or nuclear marker DAPI (5μg/mL). Midguts were mounted in Vectashield Plus antifade mounting medium.

Female flies were used for all midgut experiments because the female midgut exhibits an enhanced damage response, characterized by a higher density of ISCs, and better proliferative capacity and survival following tissue damage compared to males (Regan et al., 2016).

### Confocal image acquisition and analysis

Wing disc and midgut samples were imaged using Zeiss LSM880 and Leica SPE confocal laser scanning microscopes. Images were processed and analyzed using FIJI software. One confocal z-stack was acquired from each wing disc and two non-overlapping z-stacks from the posterior midgut of each sample. The posterior midgut was used for all imaging experiments, as this region consistently provided the most reproducible immunostaining results.

Mean fluorescence intensities of P-Akt, P-p38, and ROS were quantified using FIJI. Phospho-histone H3 (P-H3)-positive cells were manually counted using a confocal microscope.

Because P-p38 immunostaining after H₂O₂ treatment showed high variability between samples, data were analyzed using the posterior-to-anterior (P/A) fluorescence intensity ratio rather than absolute mean fluorescence intensity. Normalization of the posterior compartment signal to the anterior compartment signal within the same wing disc provided an internal control, minimizing inter-sample variability and improving the robustness of the quantitative analysis. P/A ratios were calculated by dividing the mean fluorescence intensity of the posterior compartment (experimental region) by that of the anterior compartment (internal control) for each wing disc. Posterior and anterior compartments were identified either by GFP expression in the **hedgehog** (*hh*) posterior domain or by immunostaining for Cubitus interruptus (Ci), which marks the anterior compartment. Fold change was calculated by dividing each measurement by the mean value of the control group.

### Generation of PTEN C79A and C132A Mutations

Mutantions affecting the oxidizable cysteine residues of PTEN were generated by substituting cysteine with alanine (C79A and C132A). Alanine was selected because its small, non-reactive side chain eliminates the thiol group responsible for redox-dependent and other post-translational modifications while minimizing structural and steric perturbations to the protein. This strategy allows assessment of the specific contribution of cysteine oxidation to PTEN function. Mutant constructs, as well as a wild-type form of PTEN, were cloned in a UAS-containing vector to enable inducible and tissue-specific ectopic expression.

BDGP clone IP16020 was used as template to PCR *Drosophila* PTEN cDNA with primers PTEN-BamHI-Fwd and PTEN-XhoI-Rev. The 1564bp product was cut BamHI/XhoI and cloned in pUASt BglII/XhoI to get the vector pUASt-PTEN_Wt. To generate the C79A mutation, two independent PCRs were performed using pUASt-PTEN_Wt as template. The first PCR was carried out using primers PTEN-EcoRI-Fwd and PTEN-C79A-Rev, with the mutation C79A incorporated into the reverse primer. The second PCR used primers PTEN-C79A-Fwd and PTEN-ERI-Rev, introducing the mutation on the forward primer. A third PCR was then performed on the mixed PCR1 and PCR2 products using PTEN-EcoRI-Fwd and PTEN-ERI-Rev. The final product carrying the C79A mutation was cut EcoRI and the 996bp band was cloned in the pUASt-PTEN_Wt digested and excised EcoRI fragment to get the vector pUASt-PTEN_C79A. The same strategy was used to introduce the C132A mutation using primers PTEN-EcoRI-Fwd and PTEN-C132A-Rev for the first PCR and PTEN-C132A-Fwd and PTEN-ERI-Rev for the second PCR. A third PCR was performed on the mixed PCR1 and PCR2 products using PTEN-EcoRI-Fwd and PTEN-ERI-Rev. The final product of 996bp carrying the C132A mutation was cut EcoRI and cloned in the pUASt-PTEN_Wt digested and excised EcoRI fragment to get the vector pUASt-PTEN_C132A. All constructs were verified by restriction analysis and DNA sequencing. Constructs were injected for transgenic generation in the w^1118^ background, with insertion in the second chromosome (FlyORF Drosophila Injection Service, Zurich, Switzerland).

Primer List 5’ to 3’:

PTEN-BamHI-Fwd: ACGGGATCCATGGCCAACACTATTTCGT PTEN-XhoI-Rev: ATCCCTCGAGTTACAGGTATGTTGATTCA PTEN-C79A-Fwd: ATCTATAACCTAGCATCGGAGCGTAGT PTEN-C79A-Rev: ACTACGCTCCGATGCTAGGTTATAGAT PTEN-C132A-Fwd: GTAGCCGTGCACGCAAAAGCTGGAAAG PTEN-C132A-Rev: CTTTCCAGCTTTTGCGTGCACGGCTAC PTEN-EcoRI-Fwd: TGGGAATTCGTTAACAGATCCATGGCCAACACTA PTEN-ERI-Rev: ATTGAAATCTTGAATTCTTCTGAAAATC

### GFP-Positive Cell Density Measurement

The proliferative capacity of intestinal progenitor cells was assessed by quantifying the density of GFP-positive cells in *esg>GFP* midguts. Confocal z-stacks of the posterior midgut were acquired and processed using FIJI software. GFP z-stacks were converted into maximum-intensity projections using the Maximum Intensity Projection function. The resulting images were thresholded using the Max Entropy algorithm, converted to binary, and adjacent cells were separated using the Watershed function. GFP-positive cells were quantified using the Analyze Particles tool. All segmentations were manually inspected to ensure accurate cell identification and counting. GFP-positive cell density was calculated by dividing the number of GFP-positive cells by the analyzed tissue area. To facilitate comparisons between experiments, values were normalized to the mean of the corresponding control group and expressed as fold change.

### Statistical analysis

Statistical analyses were performed using GraphPad Prism software. Unless otherwise indicated, comparisons among multiple groups were performed using one-way analysis of variance (ANOVA) followed by Tukey’s multiple-comparisons test. For the analyses presented in Fig. 5B–D, statistical significance was assessed using a two-tailed Student’s t-test. Data were considered statistically significant at *p* < 0.05. Statistical significance is indicated as follows: \**p* < 0.05, \*\**p* < 0.01, and *** *p* <0.001.

---

**Supplementary Figure 1**. Spectrum of wing phenotypes observed for the *sal^E/Pv^-LHG LexO-rpr* (*sal^E/Pv^>rpr*) *nub-Gal4* (*nub>*) experiments on Fig. 1 and Fig. 3. (A) The normal regeneration category includes wings with the complete set of veins and interveins, as well as wings lacking only a small segment of the anterior vein L2. (B) The aberrant regeneration category includes wings with partial or complete loss of one or more veins and/or intervein regions.

**Supplementary Figure 2**. Phosphatase activity is abolished in *Pten^C132A^* and preserved in *Pten^WT^* and *Pten^C79A^*. (A) Representative images of phospho-Akt (P-Akt) staining in wing imaginal discs expressing GFP, Pten^WT^, Pten^C79A^ or Pten^C132A^ for 24h in posterior compartment (*hh-Gal4*). Larvae were maintained at 17°C for 7 days and then transferred to 29°C for 24h before dissection. (B) Quantification of phospho-Akt (P-Akt) levels expressed as the posterior/anterior ratio. Compartments were identified by GFP fluorescence in the control discs (*UAS-GFP*) or by immunostaining of anti-Ci, a marker of the anterior compartment in discs expressing different transgenes (*UAS-Pten^WT^*, *UAS-Pten^C79A^* and *Pten^C132A^*). Genotypes and sample sizes are as follows: *hh>GFP* (n=16), *hh>Pten^WT^*(n=20), *hh>Pten^C79A^* (n=24) and *hh>Pten^C132A^*(n=16). Box plots show the maximum–minimum range (whiskers), the upper and lower quartiles (boxes), and the median value (horizontal line). Statistical significance was determined by one-way ANOVA followed by Tukey’s multiple-comparisons test: ns = not significant and ***p < 0.001. Images are representative of independent experiments. Scale bars 50µm.

